# Metagenomic identification of novel *Hepaciviridae* in Australian birds reveals pegivirus recombination

**DOI:** 10.64898/2026.09.17.752486

**Authors:** Jasper W. Schwarz, Josephine Rieken, Ethan Mandojana, Karrie Rose, Heather Fenton, Jane Hall, Kate Van Brussel, Edward C. Holmes

## Abstract

The *Hepaciviridae* family of RNA viruses have a wide range of hosts including arthropods, fish, birds and mammals. Birds are well documented reservoirs of these viruses, owing to their physiological and ecological characteristics that facilitates rapid transmission through populations. To expand our understanding of the avian virome, with a particular focus on the identification of novel hepaciviruses, we performed metatranscriptomic sequencing on tissues taken from Australian endemic birds such as emus, pelicans as well as migratory birds (shearwater and albatross sp., egrets, herons and cuckoos) found deceased in New South Wales, Australia between 2021 and 2025. From these data we identified two novel pegiviruses – shearwater and Australian pelican pegivirus (denoted SWPgV and APPgV) – that provided phylogenetic evidence of a putative recombination event, reflected in incongruent tree topologies. SWPgV and APPgV form a sister clade with previously described avian-associated pegiviruses in NS3 phylogenetic trees, yet fell as a unique basal group in phylogenies based on NS5B. The lack of a clear parental lineage in NS5B domains suggests that additional divergent avian-associated pegiviruses have yet to be identified. In addition, we discovered two novel orthohepaciviruses herein named short-tailed shearwater and emu orthohepacivirus (STSHpV and EHpV) that are related to other avian-associated orthohepaciviruses, indicative of a long avian association, as well as a confirmed case of pegivirus and orthohepacivirus co-infection. Combined, these avian hepacivirus findings demonstrate the importance of tissue-based viral discovery studies in uncovering novel virus species and understanding their complex evolutionary histories.

## Introduction

Australia’s geographic isolation and temperate climate allow for a wide range of zoologically distinct flora and fauna, including more than 900 unique species of bird. Endemic species such as the emu (*Dromaius novaehollandiae*) have evolved unique characteristics to aid their survival, while birds like the Australian pelican (*Pelecanus conspicillatus)* have retained key phenotypes with their sister species that inhabit other continents. Several hundred bird species migrate to Australia from Asia and the Pacific to escape the northern hemisphere winter and to breed. Indeed, it is estimated that over 18 million seabirds such as shearwaters (*Ardenna* sp.) and gannets (*Morus* sp.) travel over 15,000 kilometres each year, with exhausted and starving birds often found washed up on the shores of Japan, North America and Australia (Skira, 1990).

Birds are important reservoirs of viral infections, in part reflecting physiological and ecological characteristics that facilitate transmission. Flight allows the circulation of pathogens to new environments and hosts, while congregating in large numbers during breeding and migration enables the rapid transmission of viruses among populations. This includes at the wild bird-poultry interface, which in turn often underpins subsequent spillover events into humans (Ayala et al., 2020; Harvey & Holmes, 2022; Nabi et al., 2021; Pettersson et al., 2020; Wille & Holmes, 2020). Accordingly, migratory birds play a critical role in the introduction of novel viruses into Australia (Wille et al., 2022; Wille et al., 2019). This is exemplified by the recent detection of the H5 subtype of highly pathogenic avian influenza virus (HPAIV, *Alphainfluenzavirus influenzae*) in migratory birds, including brown skuas (*Stercorarius antarcticus*) and southern giant petrels (*Macronectes giganteus*), representing the first report of this virus in Australia (Neave et al., 2026).

Australian birds are reservoirs for several species from the *Hepaciviridae* family of RNA viruses (previously classified within the *Flaviviridae*). Hepaciviruses are small, enveloped viruses with a positive-sense, single-stranded RNA genome that infect a wide range of hosts including birds, fish, reptiles and mammals. Traditionally, hepacivirus genomes contain a 5’ and 3’ untranslated region, with a single open reading frame (ORF) that is co- and post-translationally cleaved into three structural (capsid, pre-matrix and envelope) and at least seven non-structural proteins (NS1, NS2, NS3, NS4A, NS4B and NS5A and NS5B) that are necessary for replication. Due to their critical function during replication, the NS3 and NS5B proteins are relatively well conserved among hepaciviruses and are commonly used as phylogenetic markers (Brand et al., 2017). NS3 contains two domains: a N-terminal protease domain and C-terminal helicase domain, while NS5B contains the canonical RNA-dependent RNA-polymerase (RdRp). Phylogenies based on the RdRp have been used to classify the *Hepaciviridae* into two genera: *Orthohepacivirus* and *Pegivirus* (Simmonds et al., 2025). Orthohepaciviruses are thought to have a broad host range that encompasses birds, mammals, fish and reptiles and include the important human pathogen hepatitis C virus (HCV, *Orthohepacivirus hominis*). In contrast, pegiviruses have, to date, only been detected in birds and mammals and include human pegivirus -1 (HPgV, *Pegivirus hominis*), a non-pathogenic virus infecting human lymphocytes.

Although recombination is well documented in RNA viruses, rates of recombination vary markedly among families, in part reflecting underlying patterns of genome organisation (Pérez-Losada et al., 2015; Simon-Loriere & Holmes, 2011). The *Hepaciviridae* are thought to undergo recombination at a lower frequency than many other groups of positive-sense RNA viruses, although evidence for its occurrence can be found in each of the component genera, including both HCV and HPgV (Blackard et al., 2016; Kalinina et al., 2002) as well as bovine hepacivirus (Ma et al., 2025; Xu et al., 2025).

Herein, we used a metatranscriptomic approach (i.e., total RNA sequencing) to reveal the viruses present in a wide range of tissue types from deceased Australian birds. In doing so, we describe two novel avian-associated pegiviruses with a likely recombinant phylogenetic history, as well as two novel orthohepaciviruses

## Methods

### Animal Ethics

Samples were collected by the Australian Registry of Wildlife Health at Taronga Zoo, Sydney, Australia, under scientific license number SL100104, granted by NSW Department of Climate Change, Energy, the Environment and Water under Part 2 of the Biodiversity Conservation Act 2016.

### Sample collection and processing

Birds found deceased in New South Wales (NSW), Australia, between 2021 and 2025 were reported to the NSW National Parks and Wildlife service, NSW Department of Primary Industries and Regional Development, or the Australian Registry of Wildlife Health and transported to Taranga Zoo, Sydney for a full necropsy. Carcasses were stored at 4^°^C before being dissected to remove brain, liver, lung, spleen and kidney tissue. Skin samples were also included when lesions were noted grossly. Individual case submission details and diagnoses are presented in Supplementary Table 1. Tissues were frozen in cryovials at -80^°^C for long-term storage, then defrosted briefly before a sub-sample was transferred to a new cryovial containing 1000 μL of DNA/RNA shield (Zymo Research) and stored at -80^°^C until RNA extraction.

### RNA extraction and metatranscriptomic sequencing

Samples were defrosted and aliquots transferred to individual tubes containing 600 μL of lysis buffer, 1% β-mercaptoethanol (Sigma Aldritch) and 0.5% Reagent DX (Qiagen). Samples were homogenised using the TissueRuptor II (Qiagen) before being transferred to Eppendorf tubes (Eppendorf) and centrifuged for 3 minutes at maximum speed. RNA was then extracted from the supernatant using the RNeasy Plus Mini Kit (Qiagen) in accordance with the manufacturers’ protocol. Extracted RNA was quantified on the Qubit 3.0 Fluorometer (Thermo Fisher), before checking quality and DNA contamination on the Agilent 4200 Tapestation (Agilent). Samples containing DNA were treated with the RNase-Free DNase Set (Qiagen) and cleaned up with RNeasy clean up kit (Qiagen) in accordance with manufacturers’ protocols.

Extracted RNA was pooled in roughly equimolar ratio by organ type with an average of 4.6 samples per library (minimum 1, maximum 8). This resulted in the generation of 38 sequencing libraries, including two reagent mixes which acted as a negative control library (Supplementary Table 1). Libraries were constructed using the Truseq Total RNA Library Preparation Protocol (Illumina) before ribosomal RNA (rRNA) was depleted using the Ribo Zero Plus Kit (Illumina). Samples were sequenced by paired-end sequencing (2 x 150 bp) on a NovaSeq X platform (Illumina). Library construction and sequencing was performed by the Australian Genome Research Facility (AGRF) in Melbourne, Australia.

### Identification of novel hepacivirus sequences

Read quality was assessed using FastQC (v0.12.1) (Andrews, 2023). Raw reads were trimmed using Trimmomatic (v0.35) (Bolger et al., 2014) to remove poor-quality or adaptor sequences. Trimmed reads were *de novo* assembled into contigs using MEGAHIT with default parameter settings (v1.2.9) (Li et al., 2015). Similarity-based sequence identification was performed using DIAMOND BLASTx (v2.1.10) (Buchfink et al., 2021) by comparing assembled contigs against the RdRp–scan database (May 2025) (Charon et al., 2022) with an e-value cut-off threshold of 1 × 10^−10^, and against the protein Reference Viral Database (RVDB, October 2025) (Sayers et al., 2024) with an e-value cut off threshold of 1 × 10^−4^. Hits were further compared against the NCBI non-redundant protein using DIAMOND BLASTx (Camacho et al., 2009) to discard false positives. TaxonKit (v0.18.0) (Shen & Ren, 2021) was used to assign virus taxonomy, with sequence similarity to known vertebrate viruses used to determine whether any virus identified was likely infecting bird tissue or a component of the host diet or microbiome. The ExPaSy translate tool (Gasteiger et al., 2003) was used to identify ORFs within vertebrate associated viruses which were subsequently screened against NCBI CDSEARCH (vCDD – 62456 PSSMs) to identify protein domains. If CDSEARCH failed to identify protein domains, all six reading frames were translated in Geneious Prime v2025.2.2 and further searched for protein domains using Interproscan v2.1 (Hunter et al., 2009).

### Assessment of library composition, diversity and abundance

All abundance estimates were expressed as the number of reads aligning to the selected contig per million total reads (RPM). Trimmed reads were mapped onto assembled contigs at 99% reference sequence identity using BBmap (v39.13) (Bushnell. B, 2014). Samtools (v1.6) (Li et al., 2009) was used to calculate read coverage. Abundance was then expressed as the number of reads aligned divided by the total number of trimmed reads, before being multiplied by 1 × 10^6^.

### Extension of viral contigs RT-PCR and Sanger sequencing

To extend putative viral contigs, trimmed reads were mapped onto reference sequences using BBMap at 95% similarity. Aligned reads were then manually reviewed in Geneious Prime v2025.2.2 (https://www.geneious.com) and consensus sequences extracted. RT-PCR was performed using the SuperScript IV One-Step RT-PCR (Invitrogen) on contigs that could not be extended bioinformatically. RT-PCR products were then cleaned with ExoSAp-IT and Sanger sequenced at the Ramaciotti Centre for Genomics (University of New South Wales, Sydney).

To confirm the presence of novel hepaciviruses in individual tissue samples, total RNA from positive libraries was screened using the SuperScript IV One-Step RT-PCR in accordance with the manufacturers protocol (Invitrogen). The primers used for contig extension and RT-PCR are provided in Supplementary Table 2.

### Phylogenetic analysis of the Hepaciviridae

To determine the evolutionary relationships among the novel hepaciviruses identified here, their contigs were placed in the context of the publicly available sequences using maximum likelihood (ML) phylogenetic trees inferred from amino acid alignments of the NS3 and NS5B sequence regions. Accordingly, the sequences generated here were combined with their closest relatives available on NCBI/GenBank. Sequences were aligned using MAFFT (E-INS-I algorithm) (Katoh et al., 2019). Ambiguously or poorly aligned regions were removed using TrimAl (v1.4) (Capella-Gutiérrez et al., 2009) employing the “strict” automated method. Phylogenetic trees were inferred using the ML method implemented in IQ-TREE2 (v2.3.6) (Minh et al., 2020) with the best-fit substitution model selected using ModelFinder (Kalyaanamoorthy et al., 2017). Information on sequence type, genomic region, amino acid substitution model and tree outgroups is provided in Supplementary Table 3. Node support was calculated using 1000 non-parametric bootstrap replicates and the Shimodaira-Hasegawa approximate likelihood ratio (SH-aRLT) (Guindon et al., 2010) using IQ-TREE2, with topological support set as bootstrap values ≥ 70% and SH-aRLT values ≥ 80. All data visualisation was prepared using R v4.5.0 and R packages ape, ggplot2, ggtree, phytools, tidyverse and scales. All animal silhouettes on the resulting figures were sourced from PhyloPic and animal silhouettes, labels and clade colouring were added in Adobe Illustrator 2025.

### Identification of recombinant viral sequences

Phylogenetic analysis of pegivirus NS3 and NS5B protein domains revealed differing topological arrangements indicative of recombination (see Results). To more accurately characterise such events and identify possible recombinant breakpoints, we selected representative hepacivirus sequences which were trimmed using the “gappyout*”* mode in TrimAl before being screened for recombination using the Genetic Algorithm for Recombination Detection (GARD) program (Kosakovsky Pond et al., 2006). The putative recombination breakpoints identified by GARD were then used to separate the untrimmed pegivirus and orthohepacivirus sequences into genomic regions with potentially different evolutionary histories due to recombination. An ML phylogenetic analysis of these genomic regions (re-trimmed using the *strict* mode in TrimAl) was then performed. Sequences were determined to have recombinant evolutionary histories if the phylogenies inferred either side of putative breakpoints were supported by bootstrap values ≥ 70% and SH values ≥ 80. Putative breakpoints that did not result in statistically significant recombination were considered as only potential sites of recombination.

## Results

### Overview of metatranscriptomic data

We characterised the virome of 167 tissue samples from 62 birds representing 14 species found deceased in NSW between 2021 and 2025. This resulted in the generation of 2,009,945,472 reads, with a mean read count of 52,893,302 (range 104,664 – 80,355,915) per library. *De novo* assembly of the trimmed reads resulted in the construction of 12,496,784 total contigs, with a mean of 328,862 contigs per library (range 101 - 624,781). Read and contig counts are provided in Supplementary Table 1.

### Identification of the Hepaciviridae

In total, 62 hepacivirus contigs were assembled from 10 libraries constructed from liver, kidney and lung tissue, resulting in the identification of two novel pegivirus and two novel orthohepacivirus species. In accordance with the ICTV demarcation for new species in this group (Simmonds et al., 2017), pegiviruses were determined to represent a novel species if their sequences exhibited less than 69% and 64% amino acid identity, respectively, to the most closely related NS3 and NS5B sequences. Similarly, orthohepacivirus sequences were considered novel when sequence similarities were less than 75% and 70% for NS3 and NS5B protein domains, respectively (Simmonds et al., 2017).

## Pegiviruses

### Pegivirus genome construction, confirmation and abundance

Sequences with the highest similarity to pegiviruses were detected in five libraries. Two contigs – denoted MLI2/Pegi/1 of 9,947 bp and STSLI3/Pegi/1 of 13,640 bp – were assembled from liver tissue and estimated to have a viral abundance of 34.71 and 66.34 RPM, respectively. A third contig - STSLU1/Pegi/1 - of 10,362 bp was assembled from lung tissue and estimated to have a viral abundance of 122.14 RPM. Smaller pegivirus contigs were assembled from the kidney (STSK3) and spleen tissue (IP02) libraries. Trimmed reads from STSK3 were mapped onto STSLU1/Pegi/1 at 95% reference sequence similarity. In total, 389 reads were mapped, resulting in 91.3% coverage of STSLU1/Pegi/1 and a viral abundance of 6.94 RPM (Figure 2A). The resulting genome was denoted STSK3/Pegi/1. Putative pegivirus sequences were translated into ORFs, leading to the identification of the envelope, NS2, NS3, NS4B, NS5A and NS5B proteins (Figure 2A). We were unable to characterise the p7 and NS4A protein domains in any pegivirus genome due to high sequence divergence from annotated regions. Similarly, trimmed reads from IP02 were unable to be mapped to STSLU1/Pegi/1 due to low similarity to reference sequences, such that they were instead extended using a combination of RT-PCR and Sanger sequencing. The resulting Sanger/*de novo* products were 893 and 705 bp in length with an estimated read abundance of 0.79 and 0.63 RPM and were subsequently denoted IP02/Pegi/NS3 and IP02/Pegi/NS5B, respectively (Figure 2A).

RT-PCR was performed on individual tissue samples from positive libraries using NS5B specific primers (Supplementary Table 2). MLI2/Pegi/1 was identified in a little shearwater (*Puffinus assimilis*; Registry no. 15588.6) which was found deceased on Dee Why Beach with an additional 10 birds in 2023 (Figure 1). STSLU1/Pegi/1 (Registry no. 15589.3) and STSLI3/Pegi/1 (Registry no. 15589.8) were identified in short-tailed shearwaters (*Ardenna tenuirostris*) found in poor health on Cronulla Beach with an additional eight birds in 2023. These birds were transported to Taronga Wildlife Hospital but were euthanised due to poor prognosis. STSK3/Pegi/1 was detected in a short-tailed shearwater on North Haven Beach in 2023 (Registry no. 15597.5). Collectively, these birds were all found to be cachexic, likely induced by the long seasonal migration. Two Australian pelicans (*Pelecanus conspicillatus*) from IP02 were positive for IP02/Pegi/NS3 and IP02/Pegi/NS5B. One bird (Registry no. 14649.3) was found deceased at Lake Brewster as part of a larger die-off in 2022, the cause of which remains undetermined but is suspected to be a combination of food scarcity and adverse environmental conditions. A second pelican (Registry no. 16152.1) which died from emaciation and severe nematodiasis near Port Macquarie in 2024 also retuned a positive result. No overt pathologies suggestive of viral infection were noted on necropsy or histopathology in positive individuals.

**Figure 1.**
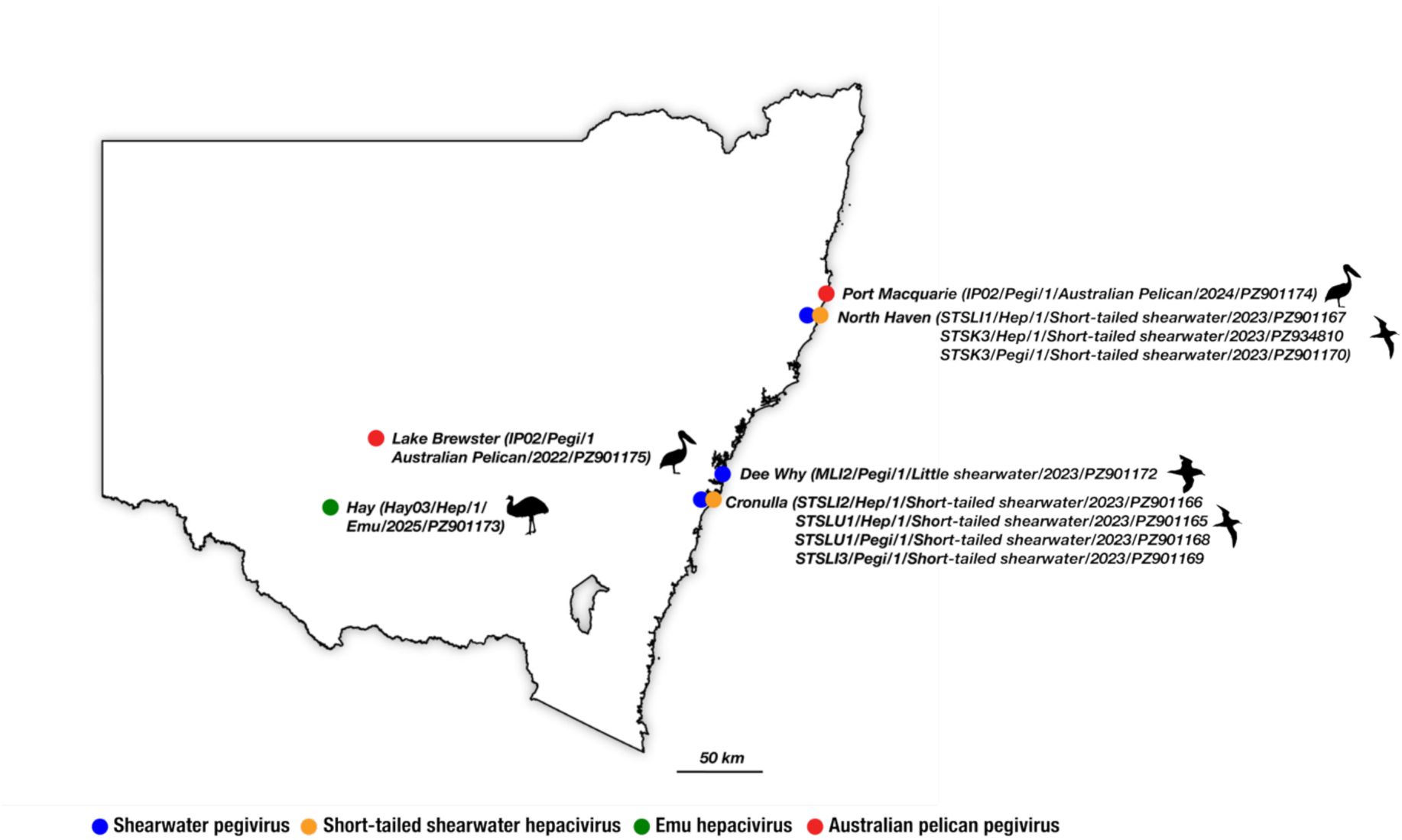
Sampling locations for avian hepaciviruses. Map of New South Wales (NSW), Australia, showing the sampling location of birds positive for hepaciviruses. The bird species from which the different viruses were sampled are coloured by blue, orange, green and red circles according to the key. Labels show contig ID/host/year/accession.

**Figure 2.**
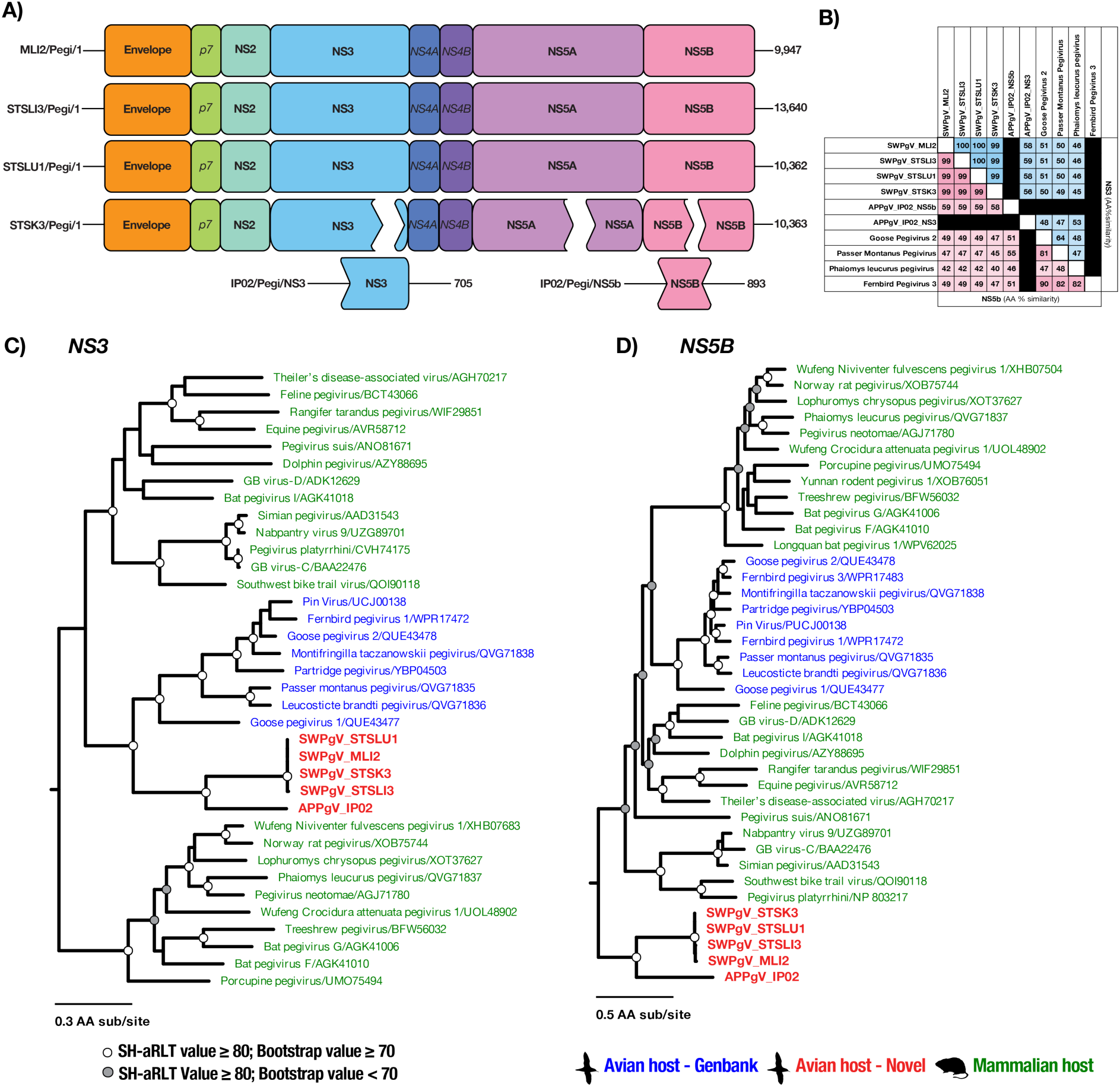
Genomic structures and phylogenetic analysis of pegiviruses. **(A)** Key protein domains for MLI2/Pegi/1, STSLI3/Pegi/1, STSLU1/Pegi/1, STSK3/Pegi/1/, IP02/Pegi/NS3 and IP02/Pegi/NS5B: envelope, NS2, NS3, NS4B, NS5A and NS5B (labelled in bold). The NS4A and p7 proteins were not characterised (labelled in italics). Unmapped regions in STSK3/Pegi/1 are represented by breaks in protein domains. Numbers on the right edge of NS5B are the lengths of assembled contigs in nucleotides. **(B)** Heatmap of SWPgV and APPgV NS5B and NS3 protein domain similarity compared to reference sequences. Cells are coloured in accordance with amino acid sequence similarity (%) between SWPgV and APPgV and their closest BLASTx hits. **(C-D)** ML phylogenetic trees were inferred using amino acid alignments of the pegivirus NS3 **(C)** and NS5B **(D)** proteins. Tip labels are coloured according to vertebrate host-class. SWPgV and APPgV are represented in bold red. Nodes with strong topological support (SH-aRLT values ≥ 80 and bootstrap values ≥ 70%) are marked with a white circle. Nodes with SH-aRLT values ≥ 80 and bootstrap values < 70% are marked with a grey circle. Branch lengths reflect amino acid substitutions per site. Trees were rooted using an orthohepacivirus outgroup.

### Evolutionary relationships and genomic structure of pegiviruses

To determine similarity to known viruses, novel pegivirus NS5B and NS3 proteins were aligned with reference hepacivirus sequences and amino acid similarities calculated (Figure 2B). As IP02/Pegi/NS3 and IP02/Pegi/NS5B were identified from the same animals, they were assumed to represent the same viral species. MLI2/Pegi/1, STSLI3/Pegi/1, STSLU1/Pegi/1 and STSK3/Pegi/1 exhibited 99% sequence similarity over NS3 and NS5B indicating these viruses are the same species. The closest known relative in the NS5B alignments was fernbird pegivirus 3 (∼49% similarity) identified in a New Zealand fernbird (*Megalurus punctatus*), while goose pegivirus 2 found in a Swan goose (*Anser cygnoides*) from China was the closest relative in NS3 alignments (∼49% similarity). In addition, pairwise comparisons revealed that these viruses shared ∼58% similarity to both regions of IP02/Pegi, indicating these viruses are likely separate species. For NS5B, the closest relative to IP02/Pegi was Passer montanus pegivirus (55%) identified in an Eurasian tree sparrow (*Passer montanus*) from China, while for NS3 the closest relative was Phaiomys leucurus pegivirus identified in a Blyth’s vole (*Phaiomys leucurus*) from China (53% sequence similarity). In both cases, NS5B and NS3 alignments meet the ICTV threshold needed for new species demarcation. As such, we propose two new species within the pegivirus genus; shearwater pegivirus (SWPgV) and Australian pelican pegivirus (APPgV) in accordance with the hosts in which they were identified.

To determine the evolutionary history of SWPgV and APPgV, ML phylogenetic trees were inferred for both the NS3 and NS5B regions. Three major clades were identified in the NS3 tree, within which viruses clustered with other lineages sampled from hosts of the same vertebrate class (Figure 2C). As a case in point, SWPgV and APPgV fell as sister lineages to one another within an avian pegivirus clade, with the remaining two clades comprising mammalian viruses. Notably, however, in the NS5B trees SWPgV and APPgV were phylogenetically distinct from the other avian-associated pegiviruses, falling as a basal clade to the pegiviruses as a whole (Figure 2D). Such topological movement is suggestive of recombination.

### Recombination among avian pegiviruses

To analyse the potential recombination among the avian pegiviruses in more detail, a subset of hepacivirus polyproteins, including SWPgV and APPgV, were screened for potential breakpoints using the Genetic Algorithm for Recombination Detection (GARD) method. This resulted in the identification of eight potential recombination breakpoints (Table 1). The pre-envelope, NS2 and NS5A regions were suspected to contain single breakpoints, while multiple breakpoints were detected in NS3 and NS5B (Figure 3). To confirm whether SWPgV and APPgV are recombinant at identified breakpoints, we next inferred ML phylogenetic trees on the genomic regions between breakpoints (labelled regions A - G) (Figure 4A-G).

**Figure 3.**
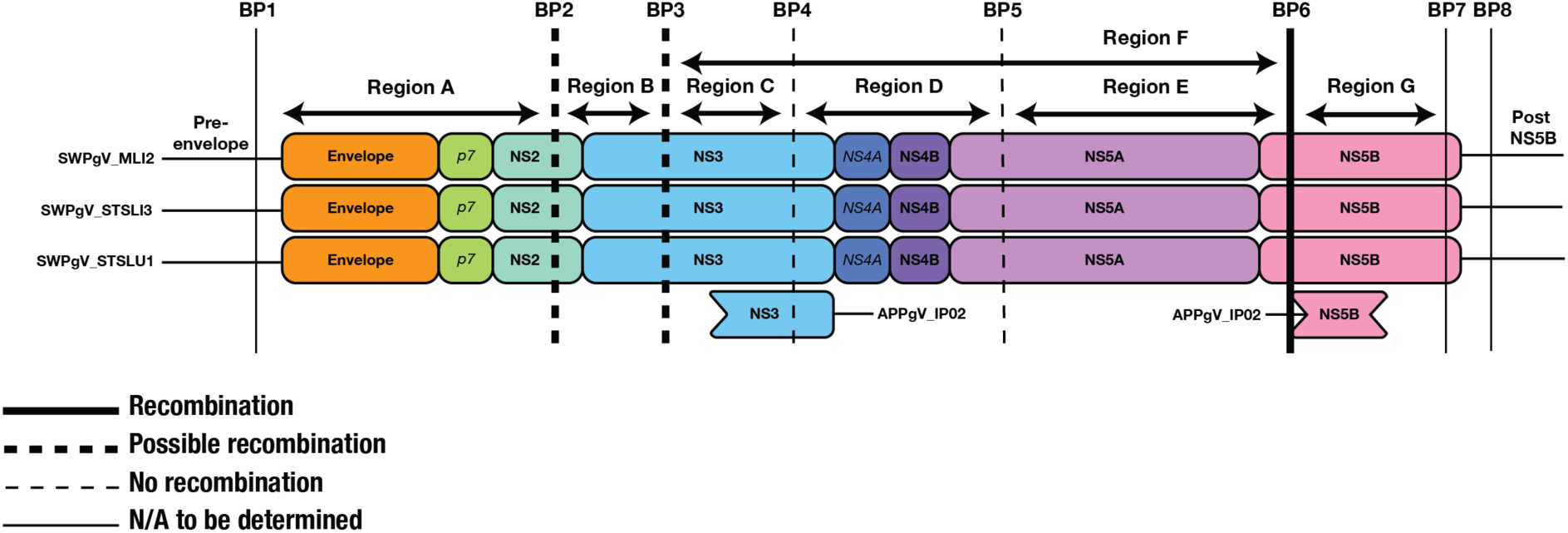
Putative recombination breakpoints and phylogenetically distinct regions within the SWPgV and APPgV genomes. Putative recombination breakpoints are displayed as vertical lines. Sites of likely recombination are marked with a solid bold line (breakpoint 6). Possible recombination breakpoints are marked with a bold, dashed line (breakpoints 2 and 3). Breakpoints with no evidence of recombination following phylogenetic analysis are marked with a thin, dashed line (breakpoints 4 and 5). Due to insufficient alignment length, breakpoints 1,7 and 8 are marked with thin, solid line and were excluded from subsequent recombination analysis. Genomic regions between breakpoints on which phylogenetic trees are inferred are marked with horizontal arrows. Specific breakpoint and region locations are provided in Supplementary Table 3.

**Figure 4.**
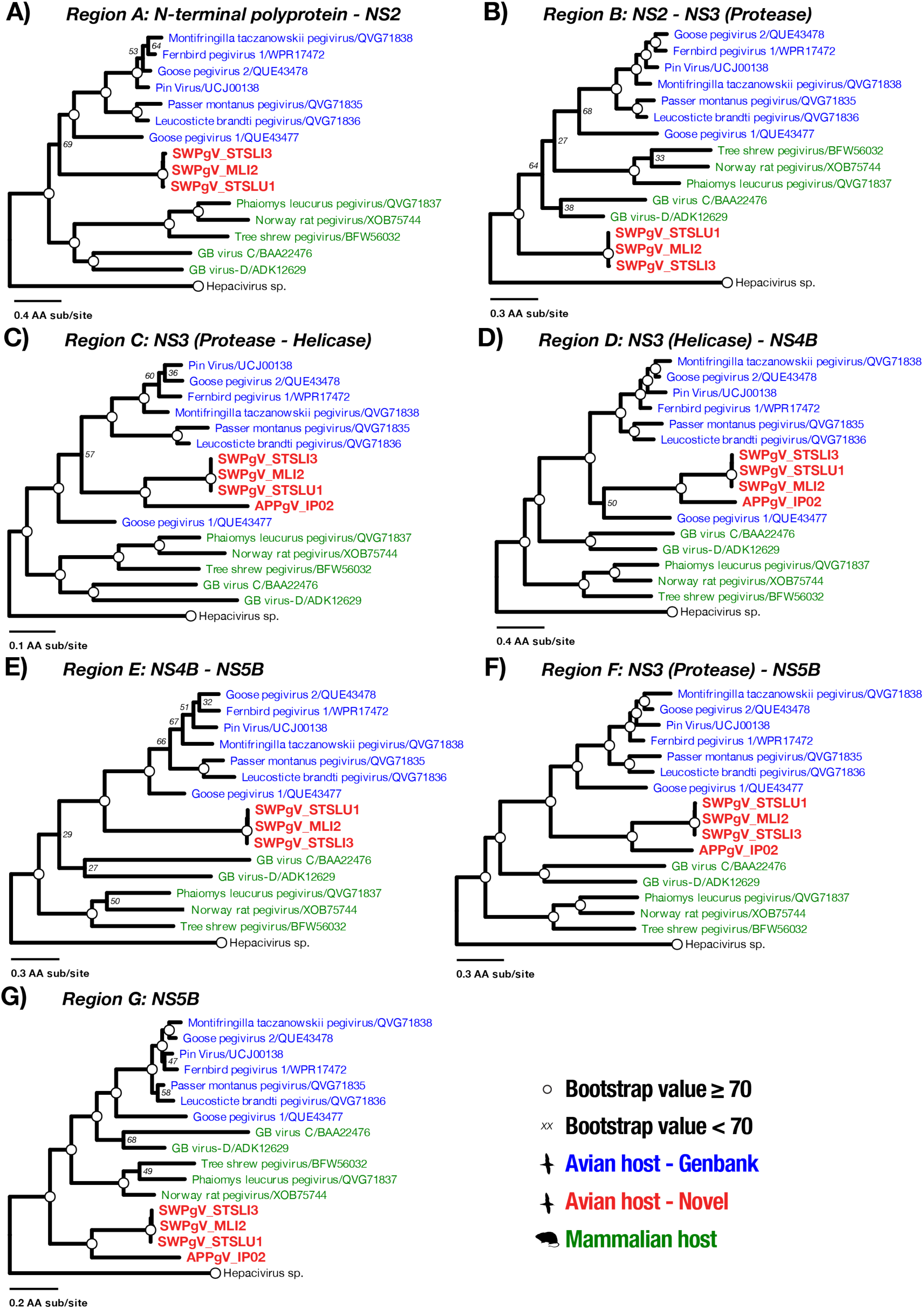
Phylogenetic analysis of putative recombinant regions in pegiviruses. ML phylogenetic trees were inferred using amino acid alignments of genomic regions between GARD-identified recombination breakpoints (Figure 3) and labelled in accordance with panel title. Tip labels are coloured in accordance with vertebrate class. SWPgV and APPgV are represented in red, bold text. Bootstrap values ≥ 70% are represented with a white circle. Bootstrap values < 70% are represented with text. Branch lengths are representative of amino acid substitutions per site in accordance with the scale. Trees are outgroup rooted on the hepacivirus sequence and are collapsed for clarity.

**Table 1.** Putative recombination breakpoints and support values identified by GARD.

| **GARD**<br>**breakpoint** | **Protein domain** | **AA**<br>**position** | **GARD**<br>**Support**<br>**(%)** | **Phylogenetic analysis**<br>**interpretation** |
| --- | --- | --- | --- | --- |
| BP1 | Pre-envelope | 61 | 99.3 | N/A |
| BP2 | NS2 | 521 | 99.3 | Possible recombination |
| BP3 | NS3 protease | 781 | 99.3 | Possible recombination |
| BP4 | NS3 helicase | 1081 | 99.3 | No recombination |
| BP5 | NS5A | 1479 | 69.5 | No recombination |
| BP6 | N-terminal NS5B | 1762 | 99.3 | Likely recombination |
| BP7 | Central NS5B | 2068 | 99.3 | N/A |
| BP8 | Post-NS5B | 2173 | 99.6 | N/A |

Trimmed alignments of regions either side of breakpoints 1 (pre-envelope), 7 and 8 (post-NS5B) were 61, 93 and 24 amino acid residues in length. As these are insufficient for reliable phylogenetic analysis we were unable to determine whether tree topology varied either side of these breakpoints. Breakpoint 2 is located within NS2. In genomic region A (i.e., upstream of breakpoint 2), SWPgV falls as a sister group to other avian pegiviruses forming a bird-associated clade (Figure 4A). This phylogenetic pattern suggests that the parent lineage of SWPgV region A was an avian-associated pegivirus. In contrast, in region B phylogenies (i.e., downstream of breakpoint 2), SWPgV does not group with the other avian-associated pegiviruses and instead falls as the basal pegivirus lineage consistent with recombination (and the overall NS5B tree – Figure 2D), although without significant bootstrap support on the key nodes (Figure 4B). As such, we suggest that breakpoint 2 is a possible site of recombination in SWPgV, although because possible recombination sites displayed low region-specific similarity to other known pegiviruses we were unable to identify parental lineages of either SWPgV or APPgV. This also suggests the existence of other, undiscovered pegivirus lineages. Breakpoint 3 is located between the protease and helicase domains in NS3. In genome region C related to this breakpoint, SWPgV once again clusters with other avian-associated pegiviruses with strong bootstrap support (although with a different topological pattern than in region A) (Figure 4C). Recombination analysis of breakpoints 2 and 3 in APPgV could not be performed as we were unable to recover regions A or B of this virus. In regions D and E (either side of breakpoints 4 and 5), the phylogenetic positioning of SWPgV and APPgV relative to goose pegivirus 1 varies, with poor topological support. Regardless, these novel pegiviruses clustered with other avian-associated pegiviruses on both sides of putative breakpoints (Figures 4D – E). As such, breakpoints 4 or 5 were not considered recombinant. Breakpoint 6 is located within the NS5B protein domain. Importantly, as earlier breakpoints (4 and 5) were not found to display signals of a recombination event, the whole genomic region between breakpoints 3 and 6 was analysed together, with an additional phylogenetic tree inferred (labelled region F) (Figure 4F). As in regions A and C - E, SWPgV and APPgV cluster with other avian-associated pegiviruses in region F. In contrast, for phylogenies inferred using region G, SWPgV and APPgV again fall as the most basal pegivirus sequences (Figure 4G). However, unlike regions A and B (breakpoints 2 and 3), topological placements are well supported in regions F and G and exhibit strong phylogenetic incongruence – indicative of recombination – compared to phylogenies in which SWPgV and APPgV cluster with the other avian viruses.

## Orthohepaciviruses

### Orthohepacivirus genome construction, confirmation and abundance

Orthohepacivirus sequences were identified in four short-tailed shearwater tissue libraries. Two orthohepacivirus genomes of length 10,137 and 10,346 bp were recovered from liver tissue library STSLI2 (denoted STSLI2/Hep/1) and lung tissue library STSLU1 (denoted STSLU1/Hep/1). Sequences had an estimated abundance of 37.11 and 108.42 RPM respectively. STSLI2/Hep/1 and STSLU1/Hep/1 were translated into ORFs and screened using Interproscan with the envelope, NS2, NS3, NS4B and NS5B identified (Figure 5A). Regions within the STSLI2/Hep/1 and STSLU1/Hep/1 polyprotein ORF where the capsid, p7, NS4A and NS5A proteins are expected be found were also present, but we were unable to accurately characterise these domains due to high levels of sequence diversity when compared to annotated reference genomes (Figure 5A). Several small orthohepacivirus contigs were also identified in kidney tissue library STSK3 (denoted STSK3/Hep/1) and liver tissue library STSLI1 (denoted STSLI1/Hep/1). Trimmed reads from libraries STSK3 and STSLI1 were mapped to STSLI2/Hep/1 and STSLU1/Hep/1 at 95% sequence identity. A total of 562 reads were mapped from STSK3, resulting in 98.5% coverage of the STSLU1/Hep/1 polyprotein, including complete NS5B and NS3 proteins (Figure 5A) and an estimated abundance of 10.02 RPM. A total of 57 trimmed reads from library STSLI1 were mapped to NS5B of library STSLU1, resulting in 64.5% coverage an estimated abundance of 1.11 RPM. No reads from STSLI1/Hep/1 could be mapped to NS3 (Figure 5). Finally, two orthohepacivirus contigs of length 6,282 and 2,314 bp were assembled from emu liver library Hay03. Contigs were joined via Sanger sequencing, resulting in an almost complete polyprotein (denoted Hay/Hep/1) with E1, NS2, NS3, NS4B and NS5B identified (Figure 5A) and estimated to have an abundance of 9.93 RPM.

**Figure 5.**
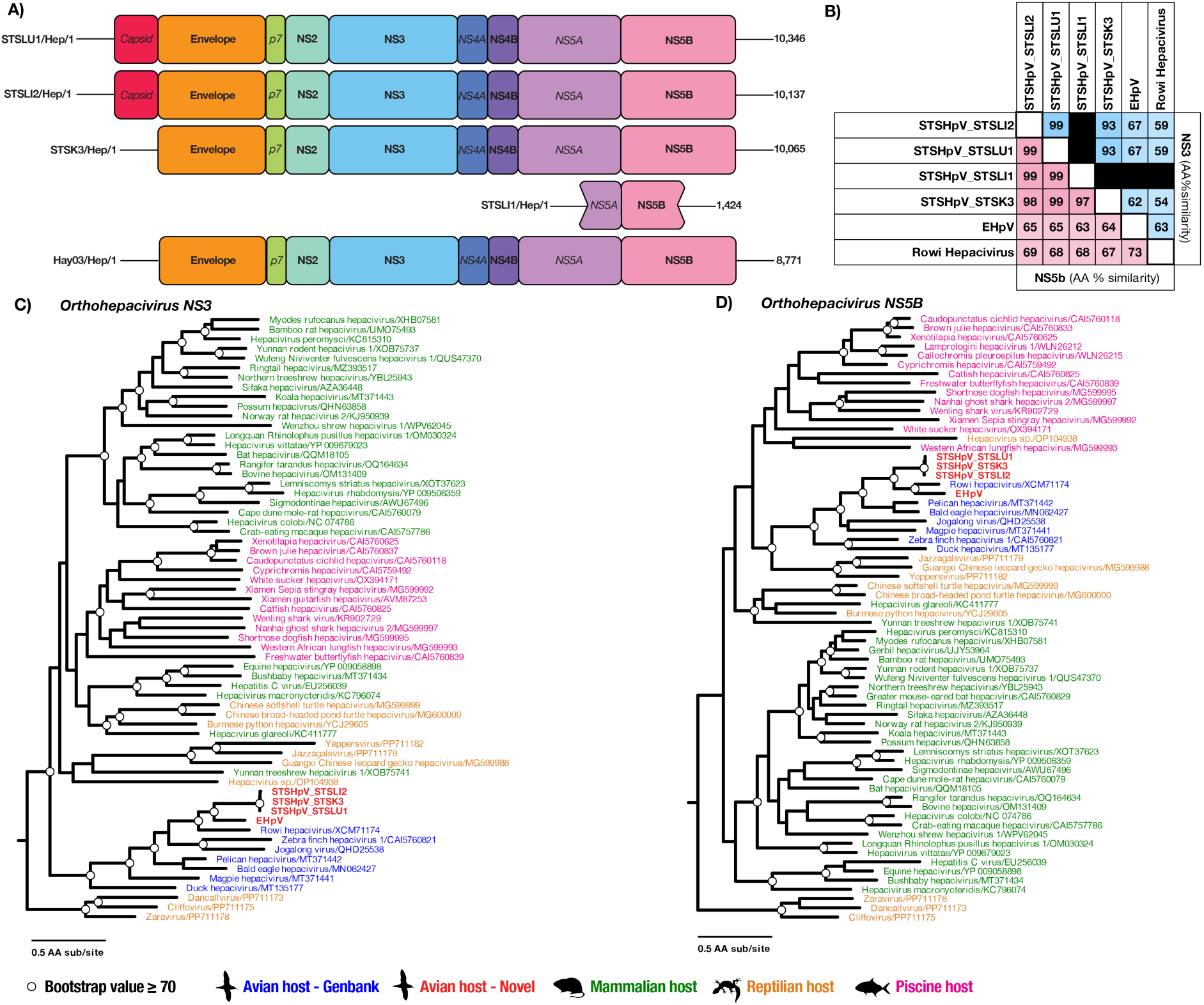
Genome structures and phylogenetic analysis of orthohepaciviruses. **(A)** Key protein domains for STSLI2/Hep/1, STSLU1/Hep/1, STSLI1/Hep/1 and STSK3/Hep/1 and Hay/Hep/1: envelope, NS2, NS3, NS4B, NS5A and NS5B identified (labelled bold). The capsid, NS4A, NS5A and p7 proteins were unable to be identified (labelled in italics). Numbers on the right edge of NS5B are the lengths of assembled contigs in nucleotides. **(B)** Heatmap of STSHpV and EHpV NS5B and NS3 protein domain similarity compared to reference sequences. Cells are coloured in accordance with amino acid similarity (%) between STSHpV and EHpV and their closest BLASTx hits. **(C-D)** ML phylogenetic trees estimated using amino acid alignments of orthohepacivirus **(C)** NS3 and **(D)** NS5B proteins. Tip labels are coloured in accordance with vertebrate class from which sequences were sampled. STSHpV and EHpV are marked in red, bold text. Nodes with bootstrap values ≥ 70% are marked with a white circle. Branch lengths are representative of amino acid substitutions per site in accordance with the scale. Trees are genus rooted for clarity.

RT-PCR was performed on individual tissue samples from all five libraries using NS5B specific primers (Supplementary Information 2). STSLI2/Hep/1was detected in 15589.6 and STSLU1/Hep/1 was detected in 15589.3. These birds were found on Cronulla beach, NSW alongside the individual described above in November 2023 (Figure 1). STSK3/Hep/1 and STSLI1/Hep/1 were detected in 15597.5 and 15597.4 which were found deceased in North Haven in 2023. Strikingly, birds 15597.5 and 15589.3 were co-infected with both STSHpV and SWPgV, representing the first confirmed case of pegivirus and orthohepacivirus co-infection of non-human hosts. Death was attributed to post-migratory cachexia. Hay/Hep/1 was present in emu 16298.5, which was euthanised in Hay, NSW due to lethargy and paralysis as part of a wider mortality event investigation. Lung and kidney tissue from this bird were also positive for this virus. No histological lesions associated with hepatitis were reported.

### Evolutionary relationships of orthohepaciviruses

To determine evolutionary relationships of putative orthohepaciviruses, a phylogenetic analysis was performed on translated NS5B and NS3 sequences and their closest relatives. Sequence similarities between novel orthohepacivirus lineages and reference sequences are presented in Figure 5B. STSLI2/Hep/1, STSLU1/Hep/1, STSLI1/Hep/1 and STSK3/Hep/1 exhibited 93% sequence similarity in NS3 and 97% similarity in NS5B, indicating that these viruses are the same species. For both proteins, rowi hepacivirus sampled from a Okarito kiwi (*Apteryx rowi*) from New Zealand was the closest reference sequence: NS3 (∼58% sequence similarity) and NS5B (∼69%). The closest relative to Hay/Hep/1 was also rowi hepacivirus for both NS3 (63%) and NS5B (73%). Pairwise comparison between shearwater-associated viruses and Hay/Hep/1 exhibit ∼67% and ∼64% sequence similarity in the NS3 and NS5B alignments, indicating that they represent two new species within the hepacivirus genus, denoted short-tailed shearwater hepacivirus (STSHpV) and emu hepacivirus (EHpV) in accordance with the hosts in which they were identified.

ML phylogenetic trees for the orthohepaciviruses were inferred using amino acid alignments of EHpV and SWPgV and reference NS3 and NS5B sequences. In a similar manner to the pegivirus phylogenies, the orthohepaciviruses broadly form clades that are associated with hosts of the same vertebrate class. Such clusters were identified in both NS3 and NS5B phylogenies, with general avian, mammalian, reptilian and piscine clades apparent. EHpV and STSHpV group with other avian-associated orthohepaciviruses in both NS3 and NS5B phylogenies. In the NS3 phylogenies, EHpV and STSHpV cluster together, with rowi hepacivirus forming the sister lineage to this group. Together, these viruses form a larger clade with jogalong virus and zebra finch hepacivirus 1, although support for this relationship is low (Figure 5C). In the NS5B phylogeny, EHpV instead clusters with rowi hepacivirus, with STSHpV forming the sister lineage to this group. This clade then groups with bald eagle and pelican hepaciviruses (Figure 5D)

## Discussion

We employed a metatranscriptomic approach to identify the viruses present in tissues from free-ranging Australian birds. From this, we identified novel pegiviruses – shearwater and Australian pelican pegiviruses – that have likely undergone interspecies recombination in at least one site in the polyprotein. In addition, the unknown nature of one of the SWPgV and APPgV parental lineages (i.e., the absence of close relatives) suggests the existence of other, currently undiscovered pegiviruses. We also describe two novel orthohepaciviruses, in a short-tailed shearwater and an emu, that fall within the avian-associated clade, as well as the first recorded example of pegivirus and orthohepacivirus co-infection in non-human hosts. Accordingly, this study highlights the importance of broad scale metagenomic testing of avian tissue for the identification of novel hepaciviruses.

Pegiviruses have been identified in a wide range of avian hosts, including Australian invasive species and the Common myna (*Acridotheres tristis*) (Chang et al., 2021). Here, we characterised the novel avian-associated pegiviruses SWPgV and APPgV. In NS3 phylogenies SWPgV and APPgV clustered with other bird-associated lineages suggesting that the parent-lineage of this genomic region has a long association with such hosts. In contrast, for phylogenies inferred using NS5B, SWPgV and APPgV fell as the most basal pegivirus lineages, separate to other avian-associated viruses, yet of unknown parental ancestry. Although such phylogenetic incongruence between conserved protein domains is indicative of a recombination event, care should be taken in drawing conclusions from these analyses due to often poor topological support reflecting high levels of sequence divergence. Genomic regions either side of breakpoints 2 and 3 had similar phylogenetic patterns to confirmed recombination breakpoint 6. This suggested additional recombination breakpoints, but which lacked strong topological support. As such, we were unable to classify either breakpoint 2 or 3 as a site of recombination.

Recombination has been previously documented in hepaciviruses but appears limited to parent lineages from the same viral species (Blackard et al., 2016; Kalinina et al., 2002; Ma et al., 2025; Twiddy & Holmes, 2003; Worobey & Holmes, 2001; Xu et al., 2025). Given the host specificity of pegiviruses (Geoghegan et al., 2017; Mifsud et al., 2023; Porter et al., 2020), it seems reasonable to assume that the parental viruses of SWPgV and APPgV must also have been capable of avian infection. However, low sequence similarity to known pegiviruses meant these species were unable to be characterised here. Hence, this strongly suggests the existence of undiscovered avian-associated pegiviruses.

Previously, mammalian-associated pegiviruses were thought to be the most basal lineages in phylogenies of this group. However, in the phylogenetic trees inferred here, the avian-associated SWPgV and APPgV fell as most basal lineages in some gene regions. Additionally, the paraphyletic nature of SWPgV and APPgV with other bird-associated pegiviruses is indicative of multiple introductions of these viruses into avian hosts.

However, as the parent lineages of SWPgV remain uncharacterised, we were unable to determine whether phylogenetic positioning is replicated in NS3. In addition, topological support in the complete NS3 and NS5B phylogenies was sometimes poor. As a consequence, it is difficult to ascertain the validity of these proposed evolutionary histories. Additional sampling of bird tissue is required to characterise undiscovered pegivirus lineages.

Orthohepaciviruses have an equally wide range of avian hosts, including ducks, kiwis and passerines (Mifsud et al., 2023; Taylor et al., 2025). Herein, we characterised two novel orthohepaciviruses sampled from bird tissue – STSHpV and EHpV. Orthohepaciviruses are broadly thought to have evolved through virus-host co-divergence, with cross-species transmission events occurring within host classes (Geoghegan et al., 2017; Hartlage et al., 2016; Mifsud et al., 2023; Porter et al., 2020; Shi et al., 2018). Indeed, the relatedness of STSHpV and EHpV to other avian orthohepaciviruses supports a long-term association with bird hosts. Furthermore, clustering of viral lineages from evolutionary-related hosts such as emus and okarito kiwis is consistent with virus-host co-divergence occurring at a deeper taxonomic level over tens of millions of years (Hartlage et al., 2016; Mifsud et al., 2023). Interestingly, STSHpV also clusters with these viruses, despite having been sampled in ecologically and phylogenetically distinct hosts. Collectively, this suggests that STSHpV and EHpV likely has an evolutionary background of long-term virus-host co-divergence, yet with more recent cross-species virus transmission. Further sampling of phenotypically similar, but evolutionarily divergent host-taxa, including cassowaries (Casuariiformes), ostriches (Struthioniformes) and rheas (Rheiformes) could allow for more accurate differentiation between evolution pathways of avian associated orthohepaciviruses (Maderspacher, 2022).

The detection of SWPgV and STSHpV in the same host represents the first recorded example of pegi- and orthohepacivirus co-infection in a non-human host. In humans, HCV infection is frequently accompanied by HPgV co-infection, likely reflecting similar transmission routes (Ng et al., 2015; Nunes et al., 2023; Pybus & Thézé, 2016; Stapleton, 2022). However, these such events are predominantly associated with blood transfusions and intravenous drug use which is not applicable in a wildlife setting. Consequently, the mechanisms which facilitate pegi and orthohepacivirus co-infection like presented here remains unclear. Indeed, these findings re-affirm a lack of understanding in how these viruses are transmitted in non-human hosts. Further investigation into such pathways is required to understand how co-infection arise and are maintained in wildlife populations.

In sum, through the metagenomic surveillance of avian tissue samples we reveal multiple novel hepaciviruses which provide greater detail into the dynamic evolutionary processes of these viruses. In doing so, we provide evidence that these viruses are able to undergo interspecies recombination previously thought to be unlikely. These analyses highlight the importance of tissue sampling in future virome studies and the opportunities for future research which would allow for a better understanding into transmission and evolutionary mechanisms of such viruses in avian populations.

## Data Availability

All the raw sequence reads generated here are available on the NCBI SRA under BioProject <u>PRJNA1519063</u> and accession numbers SAMN62711053 - SAMN62711059 and SAMN63166292. All viral genomes and corresponding sequences assembled in this study have been deposited in GenBank under accession numbers PZ901165 - PZ901175 and PZ934808 - PZ934810.

## Acknowledgements

We thank Taronga Conservation Society Australia and NSW Departments of Primary Industries and Regional Development, and Climate Change, Energy, the Environment and Water for funding state-wide wildlife health investigations, which resulted in the availability of tissue samples and metadata. We acknowledge the National Computational Infrastructure (NCI) high-performance computer cluster, GADI, for providing the computing resources used for this study.

## Supplementary Information

Supplementary Table 1. Sequencing library information with pooled species, organ and necropsy information.

Supplementary Table 2. Primers used for RT-PCR confirmation.

Supplementary Table 3. Recombination breakpoints identified by GARD.

## References

Ayala, A. J., Yabsley, M. J., & Hernandez, S. M. (2020). A review of pathogen transmission at the backyard chicken-wild bird interface. Front Vet Sci 7, 539925.

Blackard, J. T., Ma, G., Polen, C., et al. (2016). Recombination among GB virus C (GBV-C) isolates in the United States. J Gen Virol 97, 1537–1544.

Bolger, A. M., Lohse, M., & Usadel, B. (2014). Trimmomatic: A flexible trimmer for Illumina sequence data. Bioinformatics 30, 2114–2120.

Brand, C., Bisaillon, M., & Geiss, B. J. (2017). Organization of the flavivirus RNA replicase complex. Wiley Interdisciplinary Reviews: RNA, 8, e1437.

Buchfink, B., Reuter, K., & Drost, H.-G. (2021). Sensitive protein alignments at tree-of-life scale using DIAMOND. Nat Meth 18, 366–368.

Camacho, C., Coulouris, G., Avagyan, V., et al. (2009). BLAST+: Architecture and applications. BMC Bioinformatics 10, 421.

Capella-Gutiérrez, S., Silla-Martínez, J. M., & Gabaldón, T. (2009). trimAl: A tool for automated alignment trimming in large-scale phylogenetic analyses. Bioinformatics 25, 1972–1973.

Chang, W.-S., Rose, K., & Holmes, E. C. (2021). Meta-transcriptomic analysis of the virome and microbiome of the invasive Indian myna (*Acridotheres tristis*) in Australia. One Health 13, 100360.

Charon, J., Buchmann, J. P., Sadiq, S., et al. (2022). RdRp-scan: A bioinformatic resource to identify and annotate divergent RNA viruses in metagenomic sequence data. Virus Evol 8, veac082.

Gasteiger, E., Gattiker, A., Hoogland, C., et al. (2003). ExPASy: The proteomics server for in-depth protein knowledge and analysis. Nuc Acids Res 31, 3784–3788.

Geoghegan, J. L., Duchêne, S., & Holmes, E. C. (2017). Comparative analysis estimates the relative frequencies of co-divergence and cross-species transmission within viral families. PLoS Pathog 13, e1006215.

Guindon, S., Dufayard, J.-F., Lefort, V., et al. (2010). New algorithms and methods to wstimate maximum-likelihood phylogenies: assessing the performance of PhyML 3.0. Syst Biol 59, 307–321.

Hartlage, A. S., Cullen, J. M., & Kapoor, A. (2016). The strange, expanding world of animal hepaciviruses. Annu Rev Virol 3, 53–75.

Harvey, E., & Holmes, E. C. (2022). Diversity and evolution of the animal virome. Nat Rev Micro 20, 321–334.

Hunter, S., Apweiler, R., Attwood, T. K., et al. (2009). InterPro: The integrative protein signature database. Nuc Acids Res 37, D211–D215.

Kalinina, O., Norder, H., Mukomolov, S., et al. (2002). A natural intergenotypic recombinant of hepatitis C Virus identified in St. Petersburg. J Virol 76, 4034–4043.

Kalyaanamoorthy, S., Minh, B. Q., Wong, T. K. F., et al. (2017). ModelFinder: Fast model selection for accurate phylogenetic estimates. Nat Met 14, 587–589.

Katoh, K., Rozewicki, J., Yamada, K. D. (2019) MAFFT online service: multiple sequence alignment, interactive sequence choice and visualization. Brief Bioinform, 20, 1160–1166.

Kosakovsky Pond, S. L., Posada, D., Gravenor, M. B., et al. (2006). Automated phylogenetic detection of recombination using a genetic Algorithm. Mol Biol Evol 23, 1891–1901.

Li, H., Handsaker, B., Wysoker, A., Fennell, T., et al. (2009). The Sequence Alignment/Map format and SAMtools. Bioinformatics, 25, 2078–2079.

Li, D., Liu, C.-M., Luo, R., et al. (2015). MEGAHIT: An ultra-fast single-node solution for large and complex metagenomics assembly via succinct de Bruijn graph. Bioinformatics 31, 1674–1676.

Ma, J., Wei, Z., Li, L., Wang, W., et al. (2025). Detection and characterization of bovine hepacivirus in cattle and sheep from Hulunbuir, northeastern China. Front Cell Infect Micro 15, 1540849.

Maderspacher, F. (2022). Flightless birds. Curr Biol 32, R1155–R1162.

Mifsud, J. C. O., Costa, V. A., Petrone, M. E., et al. (2023). Transcriptome mining extends the host range of the *Flaviviridae* to non-bilaterians. Virus Evol 9, veac124.

Minh, B. Q., Schmidt, H. A., Chernomor, O., et al. (2020). Corrigendum to: IQ-TREE 2: New models and efficient methods for phylogenetic inference in the genomic era. Mol Biol Evol 37, 2461–2461.

Nabi, G., Wang, Y., Lü, L., et al. (2021). Bats and birds as viral reservoirs: A physiological and ecological perspective. Sci Total Environ 754, 142372.

Neave, M. J., Hair, S., Mileto, P., et al. (2026) First detected incursions of avian influenza H5N1 clade 2.3.4.4b into mainland Australia from the Southern Ocean. bioRxiv, 2026.08.08.743700.

Ng, K. T., Takebe, Y., Chook, J. B., et al. (2015). Co-infections and transmission networks of HCV, HIV-1 and HPgV among people who inject drugs. Sci Rep 5, 15198.

Nunes, P., da Cruz Coelho, E., da Silva, J., et al. (2023). Hepatitis C and human pegivirus coinfection in patients with chronic hepatitis C from the Brazilian Amazon region: Prevalence, genotypes and clinical data. Viruses 15, 1892.

Pérez-Losada, M., Palero Pastor, F., Arenas, M., et al. (2015). Recombination in viruses: Mechanisms, methods of study, and evolutionary consequences. Infect Genet Evol 30, 296–307.

Pettersson, J. H., Ellström, P., Ling, J., et al. (2020). Circumpolar diversification of the *Ixodes uriae* tick virome. PLoS Pathog 16, e1008759.

Porter, A. F., Pettersson, J. H.-O., Chang, W.-S., et al. (2020). Novel hepaci- and pegi-like viruses in native Australian wildlife and non-human primates. Virus Evol 6, veaa064.

Pybus, O. G., & Thézé, J. (2016). Hepacivirus cross-species transmission and the origins of the hepatitis C virus. Cur Opin Virol 16, 1–7.

Sayers, E. W., Beck, J., Bolton, E. E., et al. (2025). Database resources of the National Center for Biotechnology Information in 2025. Nuc Acids Res 53, D20–D29.

Shen, W., & Ren, H. (2021). TaxonKit: A practical and efficient NCBI taxonomy toolkit. J Genet Genom 48, 844–850.

Shi, M., Lin, X.-D., Chen, X., et al. (2018). The evolutionary history of vertebrate RNA viruses. Nature 556, 197–202.

Simmonds, P., Becher, P., Bukh, J., et al. (2017). ICTV virus taxonomy profile: *Flaviviridae*. J Gen Virol 98, 2–3.

Simmonds, P., Butković, A., Grove, J., et al. (2025). Taxonomic expansion and reorganization of *Flaviviridae*. Nat Micro 10, 3026–3037.

Simon-Loriere, E., & Holmes, E. C. (2011). Why do RNA viruses recombine? Nat Rev Micro 9, 617–626.

Skira, I. (1990). The short-tailed shearwater: A review of it’s biology. Australian Bird Reviews 3, *15*, 45–52.

Stapleton, J. T. (2022). Human Pegivirus Type 1: A common human virus that is beneficial in immune-mediated disease? Front Immunol 13, 887760.

Taylor, J. T., Lee, V., Dearlove, T., et al. (2025). A metagenomic investigation into *Apteryx Rowi* dermatosis identifies multiple novel viruses and a highly abundant nematode. J Wildlife Dis 61, 382–395, 314.

Twiddy, S. S., & Holmes, E. C. (2003). The extent of homologous recombination in members of the genus *Flavivirus*. J Gen Virol 84, 429–440.

Neave, M., Lynch, S. E., Kirkland, P. D., et al. (2022). Australia as a global sink for the genetic diversity of avian influenza A virus. PLoS Pathog 18, e1010150.

Wille, M., & Holmes, E. C. (2020). Wild birds as reservoirs for diverse and abundant gamma- and deltacoronaviruses. FEMS Microbiol Rev 44, 631–644.

Wille, M., Lisovski, S., Risely, A., et al. (2019). Serologic evidence of exposure to highly pathogenic avian influenza H5 viruses in migratory shorebirds, Australia. Emerg Infect Dis 25, 1903–1910.

Wingett, S. W., & Andrews, S. (2018). FastQ Screen: A tool for multi-genome mapping and quality control. F1000Research, 7, 1338.

Worobey, M., & Holmes, E. C. (2001). Homologous recombination in GB Virus C/Hepatitis G virus. Mol Biol Evol 18, 254–261.

Xu, W., Wang, W., Sui, L., et al. (2025). A novel genotype of Hepacivirus bovis identified in reindeer (*Rangifer tarandus*) in northeastern China. Front Cell and Infect Micro 15, 1646191.

